# Exacerbation of sucrose-induced visceral obesity and glucose intolerance by ovariectomy and its GLP-1–dependent amelioration by the rare sugar D-allulose

**DOI:** 10.64898/2026.01.06.697863

**Authors:** Kengo Iba, Miharu Kyo, Hirotaka Ishihara, Aki Nagao, Misaki Kawabe, Kento Ohbayashi, Toshihiko Yada, Yusaku Iwasaki

## Abstract

Estrogen deficiency after menopause promotes visceral fat accumulation and insulin resistance, thereby increasing the risk of type 2 diabetes. Although hormone replacement therapy is partially effective, its use is limited by increased risks of cardiovascular disease and breast cancer, underscoring the need for safer preventive strategies. The rare sugar D-allulose has been reported to stimulate glucagon-like peptide-1 (GLP-1), a gut hormone, secretion and to improve obesity and glucose metabolism, suggesting its potential as a novel intervention for postmenopausal metabolic dysfunction. Here, we examined whether D-allulose improves obesity and glucose intolerance in a GLP-1–dependent manner under sucrose-fed conditions, using ovariectomized (OVX) female C57BL/6J mice as a model of menopause. OVX mice, but not sucrose-fed sham mice, developed exacerbated visceral obesity and glucose intolerance in response to dietary sucrose, despite similar total energy intake. Daily oral administration of D-allulose for two weeks significantly suppressed visceral fat accumulation, improved insulin resistance, and ameliorated glucose intolerance in sucrose-fed OVX mice. These beneficial effects were markedly attenuated in GLP-1 receptor knockout mice. Taken together, we found that sucrose intake after ovariectomy exacerbates visceral obesity and glucose intolerance, and that D-allulose effectively ameliorates these metabolic abnormalities. GLP-1–stimulating dietary components such as D-allulose may represent a safe and promising preventive strategy for metabolic dysfunction associated with menopause.

## 1. Introduction

The prevalence of obesity and type 2 diabetes mellitus (T2DM) is increasing worldwide, representing a major public health challenge [1]. T2DM is particularly characterized by an age-dependent increase in prevalence, and its disease burden is expected to further rise with global population aging [2].

In women, the decline in estrogen levels after menopause alters energy homeostasis and promotes visceral fat accumulation [3] as well as insulin resistance [4]. Consistent with these experimental findings, epidemiological studies have shown that postmenopausal women exhibit a higher prevalence of obesity accompanied by increased visceral adiposity and T2DM [5,6]. Hormone replacement therapy has been developed to compensate for estrogen deficiency, however, its long-term use is limited by concerns regarding increased risks of cardiovascular disease and breast cancer [7]. Therefore, there is a strong need to develop safer and more sustainable alternative strategies to prevent or ameliorate obesity and metabolic dysfunction in postmenopausal women.

Dietary factors play a critical role in the development of T2DM, and excessive intake of free sugars is considered a major risk factor [8]. Free sugars are defined as monosaccharides and disaccharides added to foods and beverages, as well as sugars naturally present in fruit juices, syrups, and honey; typical examples include D-glucose, D-fructose, and sucrose. The World Health Organization (WHO) recommends limiting the intake of free sugars to prevent non-communicable diseases, including obesity and T2DM [9]. Although the prevalence of T2DM increases with age, it remains unclear whether and to what extent free sugar intake contributes to obesity and T2DM development specifically in postmenopausal women.

In recent years, gut hormones have attracted considerable attention for their anti-obesity and antidiabetic effects. Among them, glucagon-like peptide-1 (GLP-1), an incretin hormone, has been extensively studied. GLP-1 receptor agonists improve glycemic control by enhancing insulin secretion from pancreatic β-cells and also induce body weight loss through appetite suppression [10]. However, these pharmacological agents are associated with gastrointestinal adverse effects and weight regain after treatment discontinuation, highlighting the need for more physiological and sustainable approaches to improve glucose metabolism [11–13].

Rare sugars are a group of monosaccharides that occur only in trace amounts in nature. D-allulose, a rare sugar and the C-3 epimer of D-fructose, has recently become available for dietary use owing to the establishment of large-scale production methods [14]. GLP-1 secretion is stimulated by caloric macronutrients, however, the zero-calorie rare sugar D-allulose robustly stimulates GLP-1 secretion in mice, rats, and humans [15–17].

Furthermore, studies in rodents have demonstrated that D-allulose improves hyperphagic obesity and T2DM via activation of the GLP-1 receptor [15], and human studies have shown that D-allulose suppresses postprandial glucose excursions [18]. However, whether D-allulose exerts anti-obesity and antidiabetic effects under postmenopausal conditions characterized by visceral obesity and glucose intolerance has not been investigated.

In the present study, we used ovariectomized (OVX) female C57BL/6J mice as a model of menopause. First, we examined whether subchronic sucrose intake exacerbates visceral obesity and glucose intolerance in sham-operated and OVX mice. Next, we assessed whether subchronic administration of D-allulose ameliorates sucrose-induced visceral fat accumulation and glucose intolerance in OVX mice. Finally, we investigated the involvement of the GLP-1 receptor in these effects using GLP-1 receptor knockout mice.

## 2. Results

### 2.1 Ovariectomy induces body weight gain and glucose intolerance (Experiment 1)

Body weight and glucose metabolism were compared between ovariectomized (OVX) and sham-operated female mice (Experiment 1). OVX mice exhibited a rapid increase in body weight between postoperative Days 8 and 14, and body weight was significantly higher than that of sham-operated mice from Days 12 to 29 after surgery (Figure 1A). Although body weight in OVX mice remained relatively stable after Day 30, OVX mice continued to show a modest but significant increase in body weight compared with sham-operated mice during the late postoperative period (Days 40–57) (Figure 1A).

**Figure 1.**
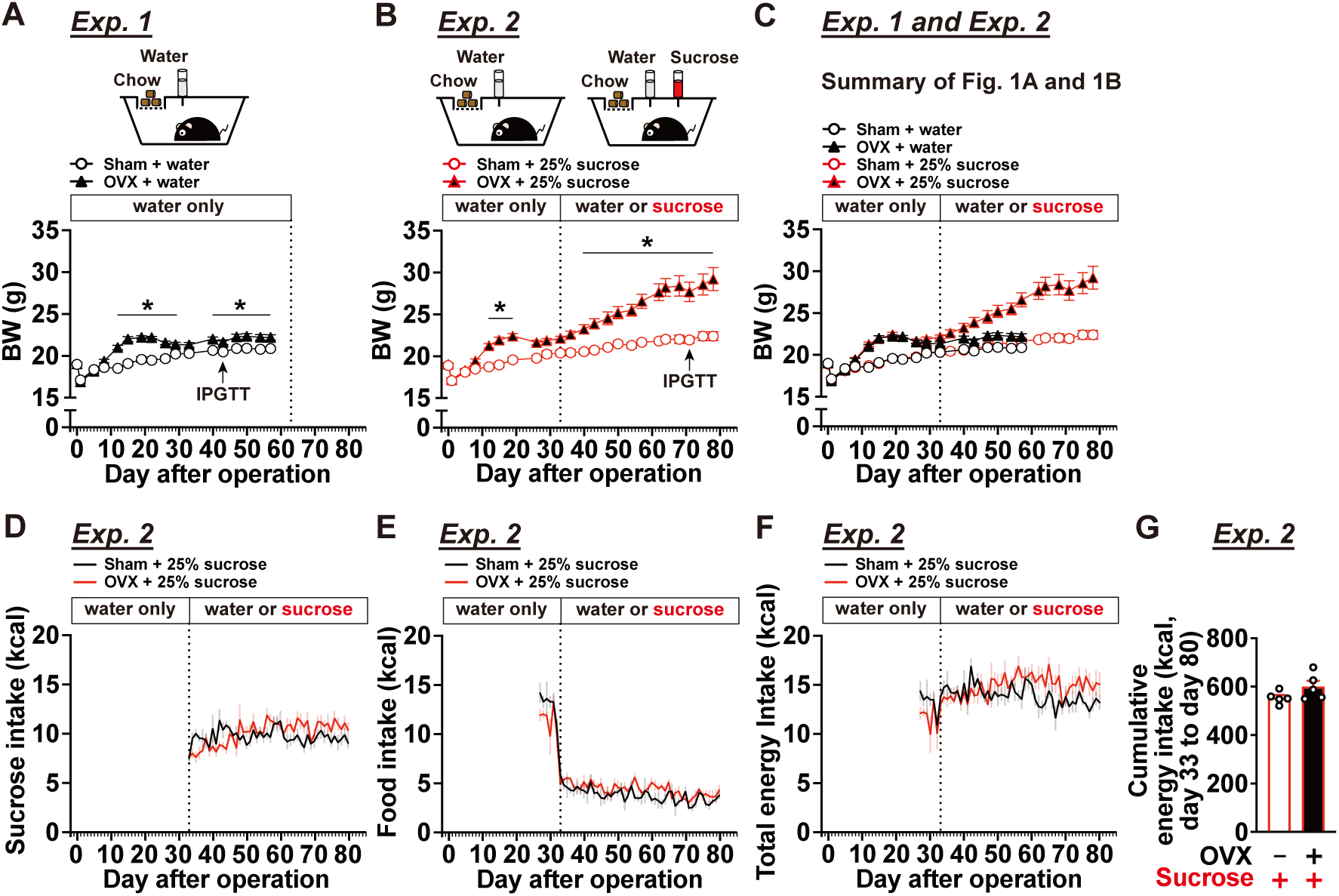
Effects of ovariectomy and two-bottle choice sucrose loading on body weight and food intake (Experiments 1 and 2) (A) Body weight changes in sham-operated or ovariectomized (OVX) mice through postoperative day 57 under a single-bottle water-only condition (Experiment 1, n = 10). (B) Body weight changes in Experiment 2 (n = 9–10); after postoperative day 32, mice were assigned to a two-bottle choice condition with 25% sucrose solution vs. water through postoperative day 78. (**C**) Integrated analysis of body weight changes from Exp.1 and Exp.2. (**D-G**) In Experiment 2, sucrose solution intake (**D**), food intake (**E**), and total energy intake calculated from sucrose and food consumption (**F**) before and after two-bottle choice paradigm, and cumulative food intake from day 33 to day 80 (**G**). n = 5. In **G**, “–” and “+” in OVX row indicate sham-operated and ovariectomized (OVX) mice, respectively. “+” in Sucrose row indicates mice provided with water containing 25% (w/v) sucrose. IPGTTs indicated in **A** and **B** denote timing of tests, and the results are shown in Figure 2. *p < 0.05 by two-way ANOVA followed by Tukey’s test for comparisons between sham and OVX.

An intraperitoneal glucose tolerance test (IPGTT) was performed on postoperative Day 43 (Figure 2A). OVX mice exhibited significantly elevated blood glucose levels at 15, 30, and 60 min after glucose administration compared with sham-operated mice (Figure 2B), resulting in a significant increase in the area under the curve (AUC) for blood glucose from 0 to 120 min (Figure 2C). In contrast, plasma insulin levels did not differ significantly between the two groups at any time point examined (Figure 2D). These findings suggest that OVX-induced glucose intolerance is associated with reduced insulin sensitivity.

**Figure 2.**
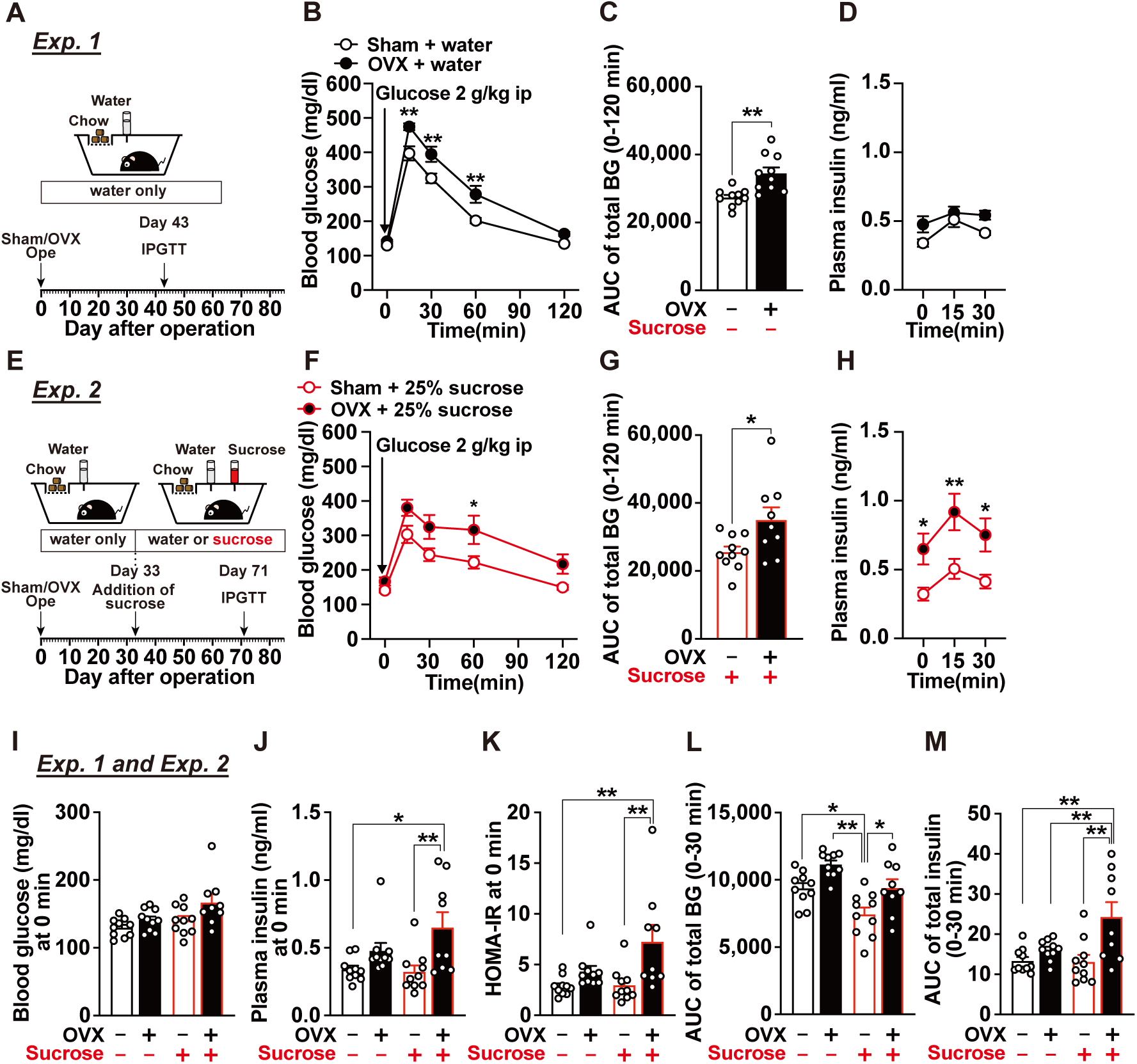
Effects of OVX and two-bottle choice sucrose loading on blood glucose levels, plasma insulin levels, and glucose tolerance (Experiments 1 and 2). (**A–H**) Experimental design of Experiments 1 (**A**) and 2 (**E**), time-course changes in blood glucose levels after intraperitoneal glucose tolerance test (IPGTT) (**B, F**) and corresponding area under the curve (AUC) (**C, G**), and time-course changes in plasma insulin levels (**D, H**). n = 9–10. In B, F, and H, *p < 0.05, **p < 0.01 by two-way ANOVA followed by Bonferroni’s test. In C and G, *p < 0.05, **p < 0.01 by unpaired t-test. (**I–M**) Integrated analyses of data from Exp.1 (**B–C**) and Exp. 2 (**F-H**). Blood glucose (**I**), plasma insulin levels (**J**), and HOMA-IR calculated from these parameters (**K**) at baseline (0 min) before IPGTT. Integrated analyses of blood glucose AUC during IPGTT derived from C and G (**L**), and AUC for plasma insulin concentrations from 0 to 30 min during IPGTT (**M**). “–” and “+” in OVX row indicate sham-operated and OVX mice, respectively, whereas “–” and “+” in Sucrose row indicate mice provided with water alone or subjected to a two-bottle choice condition with water vs. sucrose, respectively. n = 9–10. *p < 0.05, **p < 0.01 by one-way ANOVA followed by Tukey’s test.

On postoperative day 63, visceral adipose tissue mass (the sum of mesenteric, perirenal, and periuterine white adipose tissues) and fasting blood glucose levels were measured. OVX mice exhibited a marked reduction in uterine weight, confirming the effectiveness of ovariectomy (Figure 3A). Although OVX tended to increase visceral fat mass, body weight, and fasting blood glucose levels, these differences did not reach statistical significance at this time point (Figure 3B–D).

**Figure 3.**
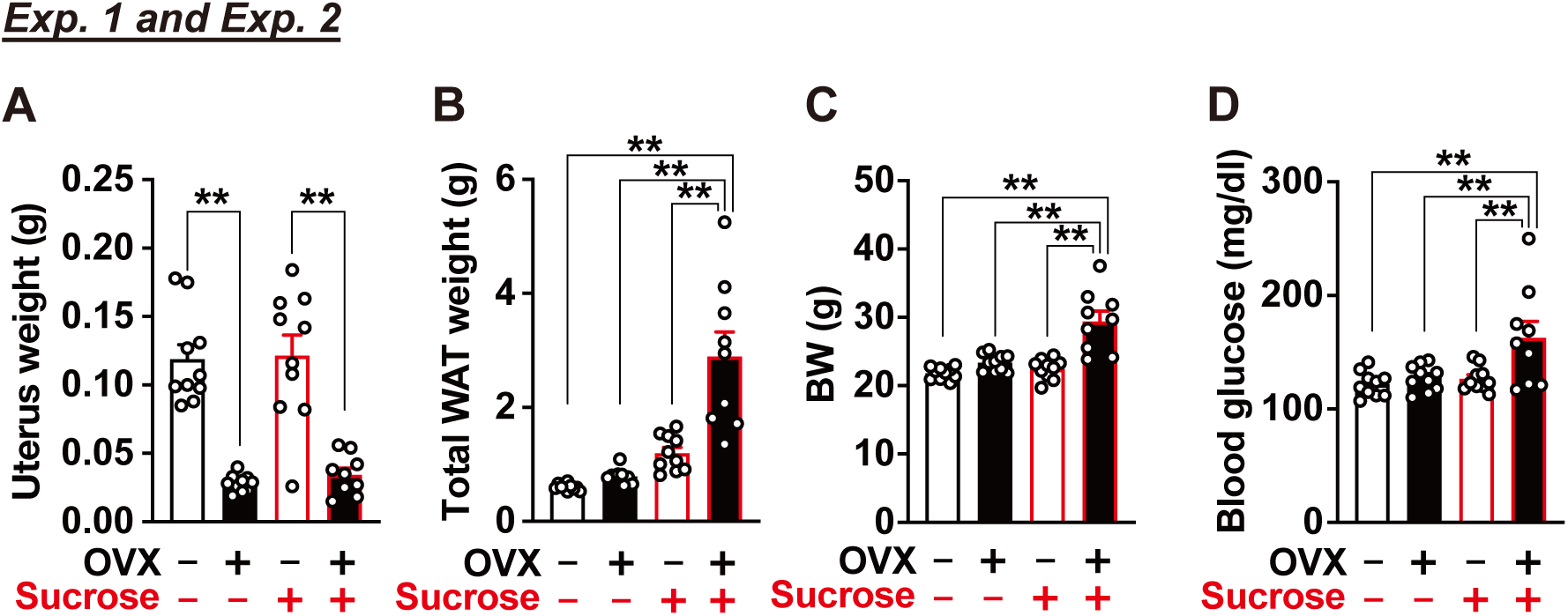
Effects of OVX and two-bottle choice sucrose loading on visceral white adipose tissue weight, body weight, and blood glucose (Experiments 1 and 2). Mice were sacrificed on postoperative day 63 in Exp. 1 and on postoperative day 82 in Exp. 2. Uterus weight (**A**), total white adipose tissue (WAT) weight including mesenteric, perirenal, and periovarian depots (**B**), body weight (BW) (**C**), and blood glucose levels (**D**) were measured. “–” and “+” in OVX row indicate sham-operated and OVX mice, respectively, whereas “–” and “+” in Sucrose row indicate mice provided with water alone or subjected to a two-bottle choice condition with water vs. sucrose, respectively. n = 9–10. *p < 0.05, **p < 0.01 by one-way ANOVA followed by Tukey’s test.

### 2.2 Excessive sucrose intake after ovariectomy exacerbates visceral obesity, glucose intolerance, and insulin resistance (Experiment 1 and 2)

Next, we examined the effects of excessive sucrose intake after ovariectomy on metabolic outcomes (Experiment 2). From postoperative day 33, when body weight gain in OVX mice had plateaued, mice were allowed free access to either water or a 25% sucrose solution using a two-bottle choice paradigm. Body weight and glucose tolerance were subsequently assessed.

In OVX mice, body weight continued to increase from Day 33 to Day 78 when sucrose was available, whereas sham-operated mice showed no marked body weight gain under the same conditions (Figure 1B). As a result, OVX mice exhibited significantly higher body weight than sham-operated mice from Days 41 to 79 (Figure 1B). Figure 1C summarizes body weight changes across Experiments 1 and 2. Notably, free access to sucrose solution did not affect body weight in sham-operated mice, whereas the same condition induced pronounced body weight gain in OVX mice.

Both OVX and sham-operated mice strongly preferred the sucrose solution (Figure 1D). Sucrose drinking reduced the intake of standard CE-2 chow in both groups (Figure 1E), however, total daily energy intake was not altered (Figure 1F). In addition, cumulative energy intake from Days 33 to 80 did not differ significantly between OVX and sham-operated mice (Figure 1G). Thus, OVX mice gained more body weight than sham-operated mice despite comparable total energy intake when sucrose was freely available.

IPGTT was performed on postoperative Day 71 (Figure 2E). OVX mice with access to sucrose exhibited significantly higher blood glucose levels after glucose loading, resulting in markedly impaired glucose tolerance (Figure 2F, G). At the same time, plasma insulin levels at 0, 15, and 30 min after glucose administration were significantly higher in sucrose-drinking OVX mice than in sucrose-drinking sham-operated mice (Figure 2H). Baseline blood glucose and plasma insulin levels at 0 min before IPGTT were compared between Experiments 1 and 2 (Figure 2I–K). OVX alone led to only modest increases in these parameters, whereas OVX mice with sucrose access showed significantly elevated fasting blood glucose and plasma insulin levels (Figure 2I, J), accompanied by a significant increase in HOMA-IR (Figure 2K). Furthermore, analysis of the 30-min AUC for total blood glucose and total insulin during IPGTT revealed that OVX alone caused only minor changes (OVX/Sucrose, –/– vs. +/–), whereas the combination of OVX and sucrose intake resulted in significant increases in both glucose and insulin AUCs (OVX/Sucrose, –/+ vs. +/+; Figure 2L, M).

On Day 82, visceral adipose tissue mass and fasting blood glucose levels were assessed. Sucrose intake did not significantly affect visceral fat mass, body weight, or fasting blood glucose levels in sham-operated mice (OVX/Sucrose, –/– vs. –/+; Figure 3B–D). In contrast, the combination of OVX and sucrose intake (OVX/Sucrose, +/+) resulted in significant increases in visceral adipose tissue mass, body weight, and fasting blood glucose levels (Figure 3B–D). Uterine weight was significantly reduced by OVX (Figure 3A).

### 2.3 Subchronic administration of the rare sugar D-allulose ameliorates ovariectomy-and sucrose-induced visceral obesity and metabolic dysfunction (Experiment 3)

We examined whether subchronic administration of the rare sugar D-allulose ameliorates metabolic dysfunction induced by OVX and sucrose intake, and whether these effects are mediated by the GLP-1 receptor using GLP-1R KO mice (Experiment 3). Female C57BL/6J mice and GLP-1R KO mice were fed a diet containing 25% sucrose and subjected to either OVX or sham surgery on Day 0. Body weight and food intake were measured daily, with cumulative food intake assessed separately during the light and dark periods at 12-h intervals (Figure 4). From Day 5 after surgery, mice received either water or D-allulose at 3 g/kg once daily by oral gavage at the onset of the light period for 2 weeks.

**Figure 4.**
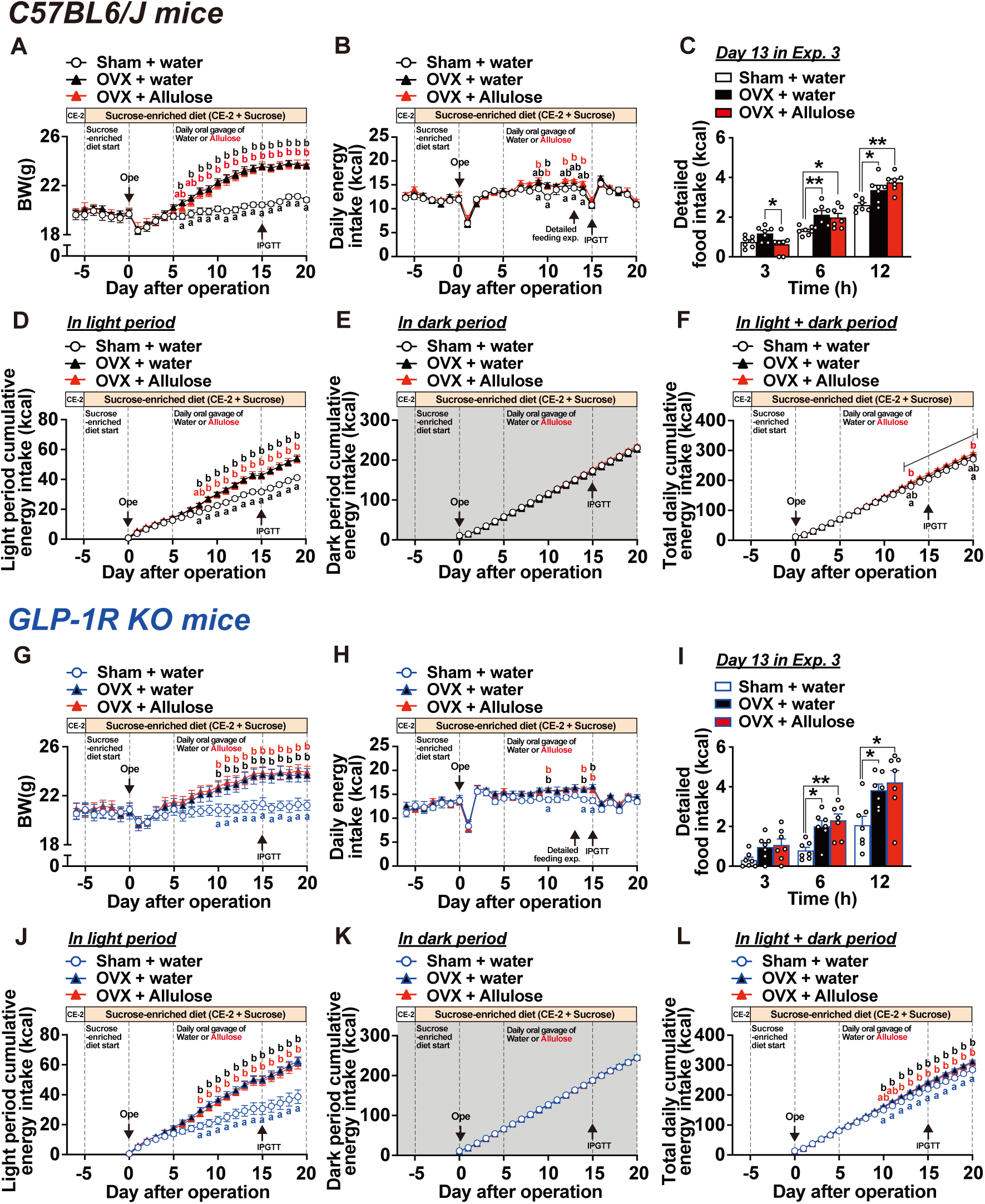
Effects of subchronic D-allulose administration on ovariectomy- and sucrose-induced light-phase hyperphagia and body weight gain and involvement of GLP-1 receptor (Experiment 3). Body weight and daily food intake were measured for 20 days after ovariectomy or sham surgery in C57BL/6J (**A-F**) and GLP-1 receptor knockout (GLP-1R KO, **G-L**) mice. From 5 days before surgery (Day −5), mice were switched from the CE-2 diet (protein, 29% of kcal) to a sucrose-enriched diet containing sucrose equivalent to 25% of total diet weight (protein, 21% of kcal). From Day 5, mice received once-daily oral gavage of water or D-allulose (Allulose) at 3 g/kg for 2 weeks at onset of light phase. Time course of body weight (**A, G**), daily energy intake (**B, H**), detailed food intake during light phase at Day 13 (**C, I**), cumulative energy intake during light period (**D, J**), cumulative energy intake during dark period (**E, K**) and total daily cumulative energy intake over whole day (24 h, **F, L**). n = 7. Different letters indicate significant differences with p < 0.05 determined by two-way ANOVA followed by Tukey’s test in A, B, D–F, G, H, and J–L. *p < 0.05, **p < 0.01 by one-way ANOVA followed by Tukey’s test in C and I.

Water-treated OVX mice exhibited a marked and continuous increase in body weight from Days 5 to 15 compared with water-treated sham-operated mice, after which body weight reached a plateau (Figure 4A). In contrast, total daily food intake in OVX mice showed only occasional increases and did not differ substantially from that in sham-operated mice (Figure 4B). Detailed analysis of food intake revealed that OVX selectively increased food intake during the light period. Light-phase food intake began to increase from Day 6, and cumulative light-phase food intake became significantly higher from Day 8 onward and continued to increase until the end of the experiment on day 20 (Figure 4D). In contrast, OVX did not affect food intake during the dark period (Figure 4E). As a result, cumulative total food intake from Days 1 to 20 showed only a modest, non-significant increase in OVX mice (Figure 4F). To determine whether OVX-induced hyperphagia during the light period contributes to body weight gain, time-restricted feeding with fasting restricted to the light period was applied (Figure S1). Under this condition, OVX-induced body weight gain was still observed (Figure S1), indicating that increased food intake during the light period plays a limited role in OVX-induced body weight gain.

Daily administration of D-allulose to OVX mice did not markedly affect body weight gain or light-phase hyperphagia compared with water-treated OVX mice (Figure 4A, B, D, and E). Unexpectedly, D-allulose slightly but significantly increased cumulative total food intake over the experimental period (Figure 4F). Although oral gavage administration of D-allulose has been reported to acutely suppress food intake [15], detailed analysis on Day 13 showed that oral gavage of D-allulose significantly reduced food intake at 3 h after administration, whereas this anorexigenic effect was no longer evident at later time points (6 and 12 h) (Figure 4C).

The OVX-induced increases in body weight and light-phase food intake observed in wild-type C57BL/6J mice were similarly observed in OVX GLP-1R KO mice (Figure 4G, H, and J–K vs. Figure 4A, B, and D–F). Moreover, continuous administration of D-allulose to OVX GLP-1R KO mice did not alter OVX-induced body weight gain or light-phase hyperphagia, consistent with the findings in wild-type mice (Figure 4G, H, and J–K). In contrast, the acute anorexigenic effect of a single intragastric administration of D-allulose was completely abolished in GLP-1R KO mice (Figure 4I vs. Figure 4C).

Next, we evaluated the effects of subchronic D-allulose administration on glucose intolerance induced by OVX and sucrose intake (Figure 5). IPGTT was performed on Day 15, after 2 weeks of D-allulose treatment. Water-treated OVX mice exhibited significantly higher blood glucose levels at 0, 15, and 30 min after glucose loading compared with water-treated sham-operated mice (Figure 5A), resulting in a significant increase in the 0–120 min AUC for total blood glucose (Figure 5B). Subchronic administration of D-allulose for 2 weeks completely reversed OVX-induced glucose intolerance, restoring blood glucose levels and glucose AUC to values comparable to those of sham-operated mice (Figure 5A, B). In contrast, the glucose-lowering effect of D-allulose was completely abolished in GLP-1R KO mice (Figure 5C, D). These results indicate that glucose intolerance induced by OVX and sucrose intake is ameliorated by sustained D-allulose administration via a GLP-1 receptor–dependent mechanism.

**Figure 5.**
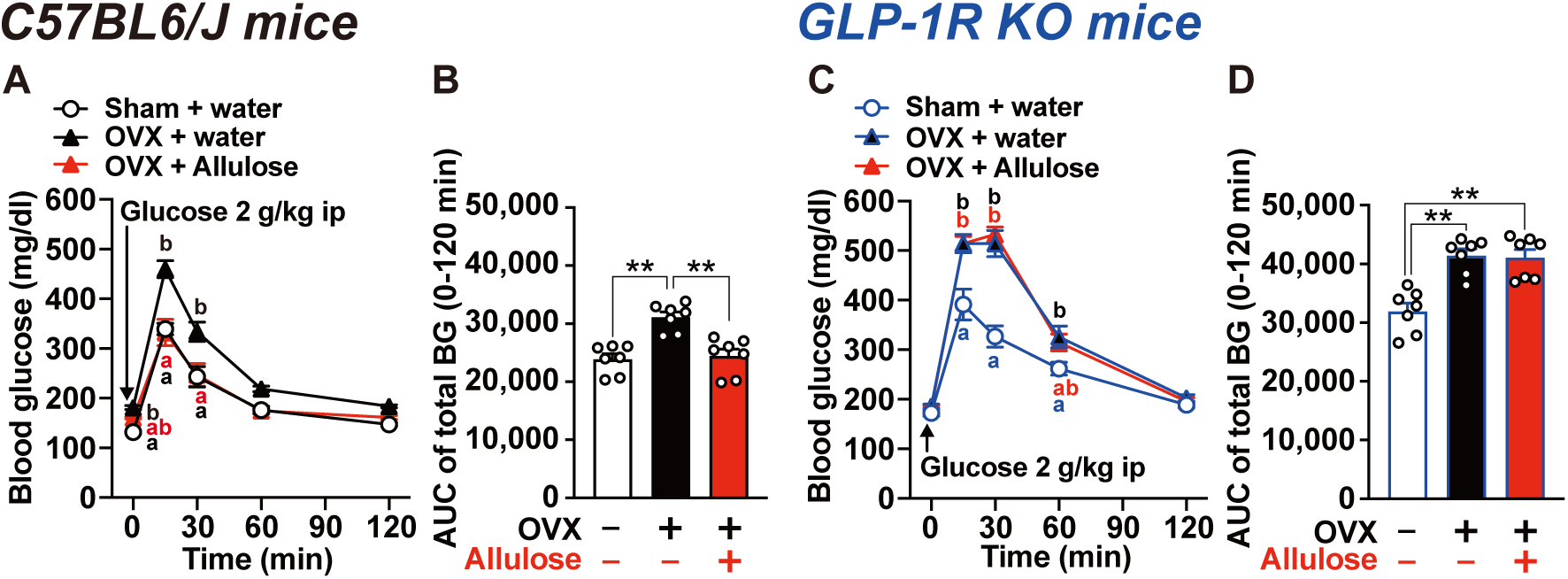
Effects of D-allulose on glucose tolerance in OVX mice fed sucrose-enriched diet and involvement of GLP-1 receptor (Experiment 3). Intraperitoneal glucose tolerance tests (IPGTTs; 2 g/kg) were performed on Day 15 of Exp. 3 in ovariectomized C57BL/6J mice (**A**, **B**) and GLP-1R KO mice (**C**, **D**). On the test day, water or D-allulose (Allulose) was not administered at onset of light phase, and IPGTTs were conducted after a 5.5-h fast. Blood glucose levels (**A**, **C**) and the AUC of blood glucose levels (**B**, **D**) during IPGTT. In B and D, “–” and “+” in OVX row indicate sham-operated and OVX mice, respectively, whereas “–” and “+” in Allulose row indicate water-treated and D-allulose-treated mice, respectively. n = 7. Different letters indicate significant differences with p < 0.05 determined by two-way ANOVA followed by Tukey’s test in A and C. **p < 0.01 by one-way ANOVA followed by Tukey’s test in B and D.

On Day 20, mice were analyzed, and a significant reduction in uterine weight was confirmed in OVX mice (Figure 6A). We then assessed weight of visceral adipose tissue, fasting blood glucose levels, and plasma insulin levels. In wild-type mice, the combination of OVX and sucrose intake significantly increased visceral adipose tissue mass, fasting blood glucose levels, plasma insulin levels, and HOMA-IR (Figure 6B, D–F). Subchronic administration of D-allulose for 2 weeks significantly attenuated these increases and restored these parameters to levels comparable to those observed in sham-operated mice (Figure 6B, D–F). In contrast, D-allulose did not markedly affect body weight gain in OVX mice (Figure 6C). In GLP-1R KO mice, OVX combined with sucrose intake significantly increased visceral adipose tissue mass, body weight, and fasting blood glucose levels, whereas plasma insulin levels and HOMA-IR were not significantly altered (Figure 6B–F). Moreover, subchronic administration of D-allulose failed to suppress the OVX- and sucrose-induced increases in visceral adipose tissue mass and fasting blood glucose levels in GLP-1R KO mice (Figure 6B, D). These findings indicate that the suppressive effects of D-allulose on visceral fat accumulation and hyperglycemia are mediated by the GLP-1 receptor.

**Figure 6.**
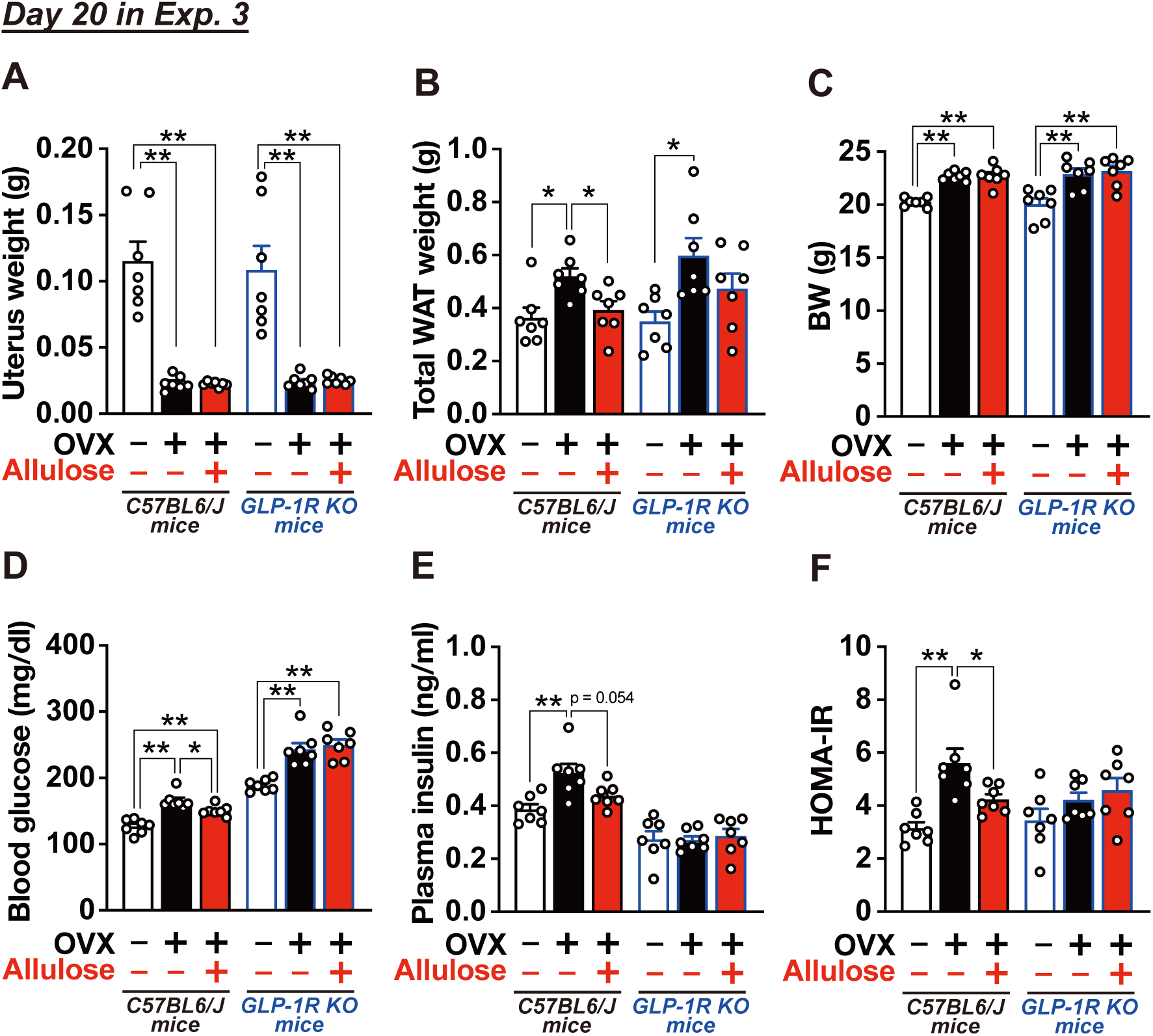
Effects of D-allulose on visceral white adipose tissue weight, body weight, blood glucose, and insulin resistance in OVX mice fed a sucrose-enriched diet (Experiment 3). Mice were sacrificed on Day 20 in Exp. 3. Uterus weight (**A**), total white adipose tissue (WAT) weight including mesenteric, perirenal, and periovarian depots (**B**), body weight (BW) (**C**), blood glucose (**D**) and plasma insulin levels, and HOMA-IR calculated from these parameters (**F**). “–” and “+” in OVX row indicate sham-operated and OVX mice, respectively, whereas “–” and “+” in Allulose row indicate water-treated and D-allulose-treated mice, respectively. n = 7. *p < 0.05, **p < 0.01 by one-way ANOVA followed by Tukey’s test, with statistical analyses performed separately within C57BL/6J and GLP-1R KO mice.

## 3. Discussion

In this study, we used OVX mice model to investigate the effects of dietary sucrose intake on metabolic homeostasis under postmenopausal conditions and to evaluate the therapeutic potential of the rare sugar D-allulose. We found that voluntary sucrose consumption markedly exacerbated visceral fat accumulation and glucose intolerance in OVX mice, despite comparable total energy intake. Importantly, subchronic oral gavage administration of D-allulose to sucrose-fed OVX mice significantly suppressed visceral adiposity and improved insulin resistance and glucose intolerance without altering total food intake or feeding rhythm. These beneficial effects of D-allulose were markedly attenuated in GLP-1R KO mice, indicating that its metabolic improvements are mediated by GLP-1 receptor–dependent signaling. Taken together, our findings suggest that metabolic dysfunction under postmenopausal conditions is strongly influenced not only by hormonal changes but also by excessive intake of free sugars, and that gut hormone–mediated pathways, particularly GLP-1 signaling, play a critical role in both disease pathogenesis and nutritional intervention.

Obesity after menopause has been attributed not to increased energy intake but to reduced energy expenditure, as demonstrated in previous studies using OVX rodent models [19]. Our results support this view, as sucrose-fed OVX mice developed overweight and visceral obesity despite similar total energy intake compared with sham-operated control mice. Estrogen regulates energy metabolism at both central and peripheral levels and promotes basal metabolic rate, physical activity, and thermogenesis in brown adipose tissue [20,21]. Therefore, loss of these estrogen-dependent mechanisms after OVX reduces energy expenditure, thereby promoting fat accumulation even without an increase in caloric intake [19]. In addition to total energy balance, accumulating evidence highlights the importance of feeding timing/rhythm in maintaining metabolic homeostasis. Independent of menopausal status, excessive food intake during the resting phase increases the risk of obesity and type 2 diabetes [22–24]. Feeding rhythms closely interact with the rhythmic expression of peripheral circadian clock genes in metabolic organs such as the liver and adipose tissue, which regulate glucose and lipid metabolism [25–28]. Estrogen deficiency induced by OVX disrupts both feeding rhythms and peripheral clock gene expression rhythms, whereas estrogen replacement largely restores these abnormalities, indicating that ovarian hormones play a key role in coordinating feeding behavior with peripheral circadian regulation [29]. In the present study, OVX mice showed increased food intake during the light phase, accompanied by body weight gain and impaired glucose tolerance. To examine whether this arrhythmic feeding directly contributes to impaired glucose metabolism, we applied a time-restricted feeding protocol that prevented food intake during the light phase. However, time-restricted feeding did not significantly improve body weight gain or glucose intolerance in OVX mice (Figure S1). Collectively, these findings demonstrate that arrhythmic feeding alone cannot fully explain the early-onset obesity and glucose metabolic abnormalities observed within 20 days after OVX.

In the present study, OVX mice given free access to sucrose in a two-bottle choice paradigm showed marked visceral fat accumulation and increased insulin resistance, whereas these changes were not observed in sham-operated mice. Importantly, although excessive sucrose intake reduced chow consumption, total energy intake did not differ significantly either before and after OVX or between OVX and sham groups. These findings indicate that the obesity and glucose metabolic abnormalities observed in this model are not attributable to increased total caloric intake. These findings further suggest that OVX enhances sensitivity to sucrose, thereby predisposing animals to obesity and glucose metabolic dysfunction. While both menopause and excessive intake of free sugars have been widely reported as independent risk factors for obesity and metabolic syndrome, animal studies directly comparing metabolic responses under identical high-sucrose conditions in the presence or absence of OVX remain limited. However, it remains unclear how estrogen deficiency following OVX alters sucrose metabolism to promote visceral fat accumulation and impair glucose homeostasis.

Sucrose is a disaccharide composed of D-glucose and D-fructose; while D-glucose functions as a major energy source for peripheral tissues, D-fructose is primarily metabolized in the liver and readily utilized for lipid synthesis [30–32]. Estrogen normally restrains lipid synthesis and regulates lipid distribution in the liver and adipose tissues [33,34]. In contrast, estrogen deficiency following OVX may weaken these regulatory controls, thereby facilitating hepatic conversion of sucrose-derived carbohydrates into lipids and their preferential accumulation as visceral fat [35]. Furthermore, accumulation of visceral adipose tissue is well known to induce insulin resistance through increased production of inflammatory cytokines and dysregulated adipokine secretion [36,37]. Consistent with these observations, sucrose-fed OVX mice in the present study showed a significant increase in visceral fat accumulation, which was accompanied by a significant elevation in HOMA-IR. These results suggest that the progression of insulin resistance associated with increased visceral adiposity is likely a major factor contributing to the worsening of glucose intolerance in this model. Taken together, estrogen deficiency induced by OVX may alter D-fructose metabolism, thereby exacerbating visceral obesity and insulin resistance. Nevertheless, this interpretation remains speculative and warrants careful consideration. Excessive sucrose intake markedly reduced normal chow consumption to approximately one-third of usual levels, resulting in decreased intake of protein, lipids, vitamins, and minerals. The potential impact of these nutritional changes on the observed metabolic phenotypes should be addressed in future studies.

In the present study, we demonstrate that subchronic administration of D-allulose ameliorates OVX- and sucrose-induced visceral obesity, insulin resistance, and impaired glucose tolerance, and that these beneficial effects are largely dependent on GLP-1R signaling. Notably, while D-allulose did not markedly suppress overall body weight gain, it selectively reduced visceral adiposity and improved glucose metabolism, suggesting a preferential attenuation of sucrose-associated metabolic dysfunction. Previous studies have shown that estrogen regulates GLP-1 synthesis and secretion in the gut and pancreas, and that circulating GLP-1 levels following glucose challenge are reduced in OVX mice but restored by estrogen replacement [38]. In addition, the anorexigenic and weight-reducing effects of GLP-1 receptor agonists have been reported to vary across the estrous cycle in accordance with fluctuations in circulating estrogen levels [39]. Together, these findings suggest that GLP-1/GLP-1R–mediated control of glucose metabolism may be compromised under conditions of estrogen deficiency. Consistent with this notion, our results show that D-allulose, which stimulates endogenous GLP-1 secretion and activates GLP-1 receptor signaling, effectively improves glucose metabolic abnormalities in OVX mice. Moreover, D-allulose has been reported to enhance energy expenditure [40,41], raising the possibility that restoration of reduced energy expenditure contributes to the observed improvements in glucose homeostasis and visceral adiposity in OVX mice. Collectively, our findings provide a mechanistic basis for the use of GLP-1–stimulating functional ingredients such as D-allulose as a safe and effective nutritional strategy to mitigate obesity and glucose metabolic dysfunction associated with menopause.

## 4. Materials and Methods

### 4.1. Mice

C57BL/6J female mice (The Jackson Laboratory Japan, Inc., Yokohama, Japan) and whole-body GLP-1 receptor knockout (*Glp1r-/-*, GLP-1R KO) mice on the C57BL/6J background generated as described previouslywere kindly provided by Dr. Daniel J Drucker (Lunenfeld Tanenbaum Research Institute, Mt. Sinai Hospital, Toronto, Canada) [42]. All mice were housed under controlled temperature (22.5 ± 2°C), humidity (55 ± 10%), and a 12-h light/dark cycle (lights on 07:30–19:30). Standard laboratory chow (CE-2, CLEA Japan, Tokyo, Japan) and water were available *ad libitum*. Purchased mice were allowed to acclimate to the facility for at least one week before any procedures. All mice aged 8-20 weeks were adequately habituated to handling before the experiments. The animal experiments were carried out after receiving approval from the Institutional Animal Experiment Committee of the Kyoto Prefectural University and in accordance with the Institutional Regulations for Animal Experiments (approval number: KPU060327-RC-4, KPU060327-RC-5).

### 4.2 Ovariectomy

Ovariectomy (OVX) was performed under anesthesia induced by intraperitoneal (IP) of a three-drug combination (MMB anesthetic) consisting of medetomidine (0.75 mg/kg; Nippon Zenyaku Kogyo, Koriyama, Japan), midazolam (4.0 mg/kg; Maruishi Pharmaceutical, Osaka, Japan), and butorphanol (5.0 mg/kg; Meiji Seika Pharma, Tokyo, Japan). Small bilateral flank incisions (∼3–4 mm) were made to identify the ovaries under a microscope, and the ovaries were surgically removed. In sham-operated mice, the ovaries were identified but not removed. The muscle layer was closed with sutures, and the skin was closed using surgical clips. After surgery, atipamezole (0.75 mg/kg, IP; Nippon Zenyaku Kogyo) was administered to reverse anesthesia, and mice were kept on a heating pad maintained at 38°C until fully ambulatory. Successful ovariectomy was confirmed by measuring plasma 17β-estradiol levels using an ELISA kit (ADI-900-174, Enzo Life Science, NY, USA) on postoperative day 26, which were significantly lower in OVX mice than in sham-operated mice (means ± SEM; 47.2 ± 5.60 pg/ml vs. 23.2 ± 6.52 pg/mL, n =5–6, p < 0.05 by unpaired *t*-test).

### 4.3 Preparation of sucrose-enriched CE-2 diet

A sucrose-enriched CE-2 diet was prepared by adding sucrose at 25% (w/w) to powdered standard CE-2 chow (CLEA Japan). In the standard CE-2 diet, protein and fat contributed 29.1% and 12.5% of kcal, respectively, whereas in the sucrose-enriched CE-2 diet, protein and fat contributed 21.0% and 4.0% of kcal, comparable to the protein content of standard laboratory diets such as AIN-93G. In contrast, the sucrose-enriched diet contained 27.9% of kcal as sucrose. Preference for the sucrose-enriched CE-2 diet vs. the standard CE-2 diet was assessed using a 48-h two-choice preference test with two food feeders. The sucrose-enriched CE-2 diet showed a preference ratio of 96.9 ± 0.83% (n = 12), indicating that it was highly palatable.

### 4.4 Experiment 1: Assessment of the effects of ovariectomy

OVX mice (n = 10) and sham-operated mice (n = 10) were housed with a*d libitum* access to solid CE-2 chow and water provided in a single bottle, and body weight was monitored until postoperative day 57 (Day 57). An intraperitoneal glucose tolerance test (IPGTT, glucose 2 g/kg) was performed on postoperative Day 43. Mice were sacrificed on postoperative Day 63. On the day of sacrifice, mice were fasted from 7:30, and blood glucose levels were measured from tail vein blood at 13:00. Mice were then euthanized by isoflurane overdose, and visceral while adipose tissue (mesenteric, perirenal, and periovarian depots) and uterus were excised and weighed. Experiments were conducted using both group housing (5 mice per cage) and individual housing. As no differences were observed in any measured parameters between housing conditions, data from both housing conditions were pooled for analysis.

### 4.5 Experiment 2: Assessment of the effects ovariectomy and two-bottle choice sucrose loading

OVX mice (n = 9) and sham-operated mice (n = 10) were housed in either group cages or individual cages with *ad libitum* access to solid CE-2 chow and water provided in a single bottle, and body weight was monitored until postoperative day 32 (Day 32). From Day 33, drinking water was provided under a two-bottle choice condition, allowing free choice between water and 25% sucrose solution. CE-2 chow continued to be provided *ad libitum* throughout this period. In individually housed mice, daily CE-2 food intake and sucrose solution intake were measured. To accurately measure fluid intake, drinking bottles equipped with a ball-bearing sipper tube to prevent leakage were used (TD-101, Tokiwa Kagaku Kikai, Tokyo, Japan). IPGTT was performed on Day 71, and mice were sacrificed on Day 82 using same procedures as in Experiment 1. Consistent with Experiment 1, housing conditions did not affect any measured parameters, and data were pooled for analysis.

### 4.6 Experiment 3: Evaluation of D-allulose effects on metabolic dysfunction induced by ovariectomy and sucrose-enriched diet

Female C57BL/6J mice or GLP-1R KO mice were used. From 5 days before surgery (Day −5), mice were switched from a standard CE-2 diet to a sucrose-enriched CE-2 diet containing 25% (w/w) sucrose and were allowed *ad libitum* access to the diet. D-allulose was provided by Matsutani Chemical Industry Co. Ltd (Itami, Japan), with purities exceeding 98%. From Day 5, mice received once-daily oral gavage of D-allulose (3 g/10 ml/kg) or water (10 ml/kg) for 2 weeks at 7:30. During the experimental period, food intake was measured at 7:30 and 19:30, and body weight was measured at 7:30. On Day 13, to assess light-phase feeding behavior in greater detail, food intake was additionally measured at 10:30 and 13:30. IPGTT (glucose 2 g/kg) was performed on Day 15. On Day 20, mice were sacrificed using the same procedures as in Experiment 1. Tail vein blood was collected at 13:00 for insulin measurements.

### 4.7 Measurements of food intake

Mice were housed and habituated for at least one week to a powdered standard CE-2 chow diet (CLEA Japan) in a feeding box (Shinano Manufacturing Co., Ltd., Tokyo, Japan). The weights of the feeding box containing the powdered food and the food spillage on the cage floor were measured at 07:30 and 19:30. Food intake was expressed in kilocalories (kcal) and calculated using an energy density of 3.4 kcal/g for the standard CE-2 chow and 3.59 kcal/g for the sucrose-enriched CE-2 diet. The intake of 25% sucrose solution was determined from changes in bottle weight, and sucrose intake was calculated using an energy density of 4 kcal/g.

### 4.8 Glucose tolerance test

Mice were fasted for 5–5.5 h (from 9:00–14:00 in Exp.1 and 2, from 7:30–13:00 in Exp. 3). Baseline blood glucose levels were measured from the tail vein, followed immediately by intraperitoneal injection of glucose at 2 g/kg. Subsequently, blood samples were collected from the tail vein at 15, 30, 60, and 120 min. Blood glucose levels were measured using the GlucoCard Plus Care (Arkray, Kyoto, Japan), and blood samples were collected using heparinized capillary glass tubes for plasma insulin assays. Plasma was collected after centrifugation (4,000 rpm, 10 min at 4°C) and stored at ‒80°C until analysis. Plasma insulin concentrations were determined using an insulin ELISA kit (Morinaga BioScience, Yokohama, Kanagawa). No allulose or water was administered on the day of the IPGTT.

### 4.9 Statistical Analysis

All data are shown as means ± SEM. Statistical analysis was performed using a two-tailed unpaired *t*-test, one-way ANOVA, or two-way ANOVA, as appropriate. When one-way or two-way ANOVA revealed significant differences, including significant main effects and/or time × factor interactions in two-way ANOVA, post hoc comparisons were conducted using Tukey’s or Bonferroni’s test. All analyses were conducted using Prism 10 (GraphPad Software, San Diego, CA, USA), and p < 0.05 was considered.

## 5. Patents

Not applicable

## Supplementary Materials

Not applicable

## Author Contributions

K.I. and Y.I. developed concept and designed the study. K.I., M.K. (Miharu Kyo), H.I., A.N., M.K. (Misaki Kawabe), K.O. and Y.I. performed experiments and analyzed data. D.J.D. developed and provided the key genetically modified mouse strain. K.I., T.Y. and Y.I. prepared figures, interpreted the results of the experiments, and drafted the manuscript. All of the authors edited the final draft. All authors have read and agreed to the published version of the manuscript.

## Institutional Review Board Statement

All animal experiments were carried out after receiving approval from the Institutional Animal Experiment Committee of the Kyoto Prefectural University and in accordance with the Institutional Regulations for Animal Experiments (Approval code and date: KPU060327-RC-4 and May 27, 2024; KPU060327-RC-5 and March 27, 2024).

## Informed Consent Statement

Not applicable.

## Data Availability Statement

The original contributions presented in this study are included in the article. Further inquiries can be directed to the corresponding authors.

## Acknowledgments

The authors thank Dr. Yuko Kawabata (Section of Oral Neuroscience, Graduate School of Dental Science, Kyushu University) for her guidance on the ovariectomy procedure, and Dr. Daniel J. Drucker (Lunenfeld-Tanenbaum Research Institute, Mt. Sinai Hospital, University of Toronto, Toronto, Canada) for providing the GLP-1 receptor knockout mice. We also thank all laboratory members including Yudai Sugiyama and Wataru Omi (Kyoto Prefectural University) for their experimental support.

## Supplemental figure and figure legends

**Figure S1.**
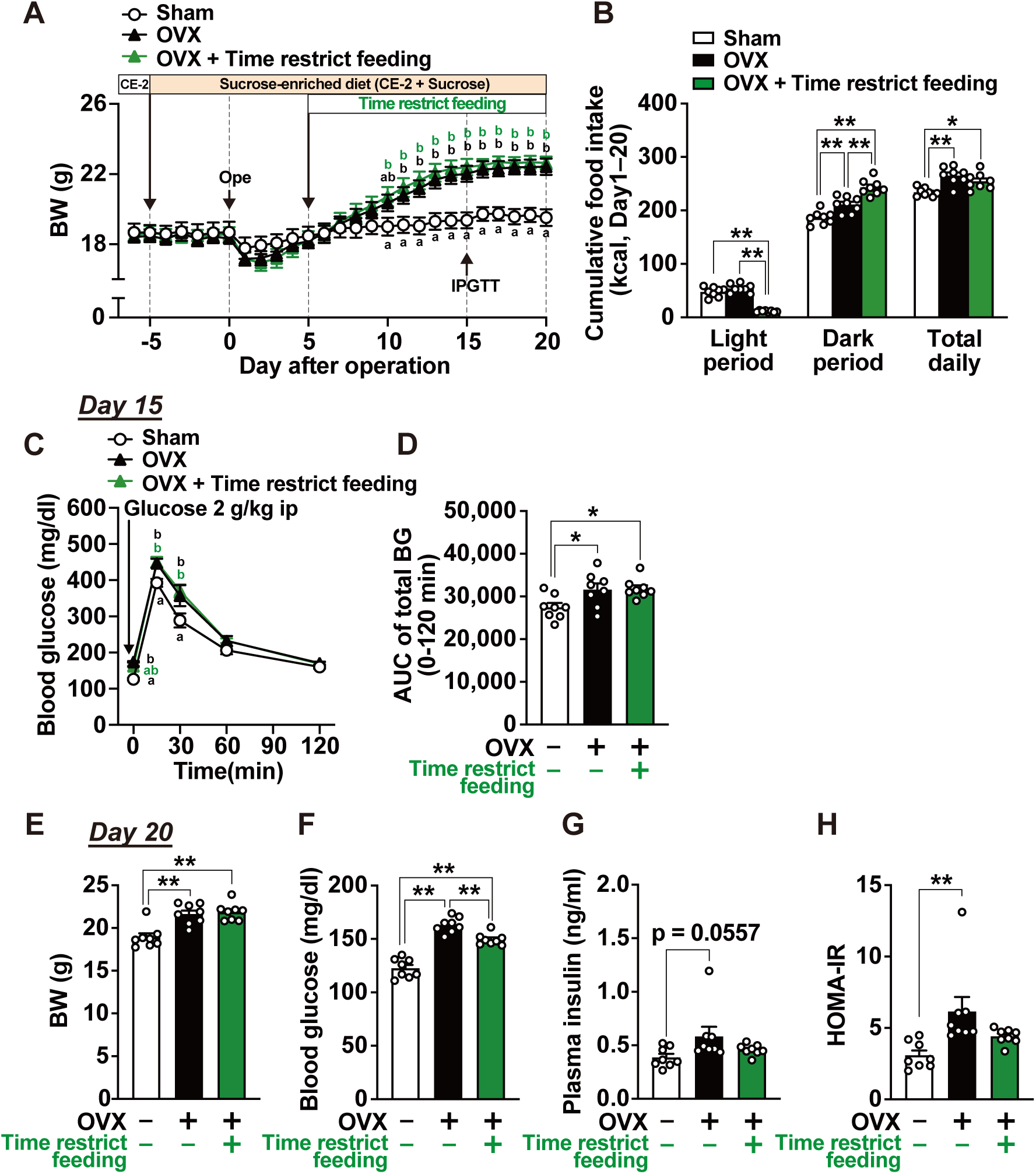
Effects of light-phase fasting from Day 5 on ovariectomy- and sucrose-induced light-phase hyperphagia and body weight gain (Experiment 4). Body weight and daily food intake were monitored for 20 days after ovariectomy or sham surgery in C57BL/6J mice. From 5 days before surgery (Day −5), mice were switched to a sucrose-enriched diet containing sucrose equivalent to 25% of the total diet weight, as in Experiment 3. From Day 5, the time-restricted feeding group was subjected to light-phase fasting. Time course of body weight (**A**); cumulative food intake during the light phase, dark phase, and total period from Day 1 to Day 20 (**B**); time course of blood glucose levels during the IPGTT and the AUC for blood glucose from 0 to 120 min on Day 15 (**C**, **D**); body weight, blood glucose, plasma insulin levels, and HOMA-IR on Day 20 (**E–H**). n = 8. Different letters indicate significant differences (p < 0.05) determined by two-way ANOVA followed by Tukey’s post hoc test in **A** and **C**. *p < 0.05, **p < 0.01 by one-way ANOVA followed by Tukey’s post hoc test in **B**, **D**, and **E–H**.

